# Knowledge and attitude regarding stem cell research and its application among medical students in Pakistan

**DOI:** 10.1101/2020.08.24.264838

**Authors:** Mashal Daud, Zaina Sajid, Tooba Ali

**Author notes:** Corresponding author: (MD). These authors contributed equally to this work.

## Abstract

**Background:** The utilization of stem cells (SCs) has led the way into a new era of therapeutics known as regenerative medicine. Their renewal property offers exciting possibilities in reversing tissue damage caused by metabolic and degenerative changes. Research should be conducted increasingly to explore the possibilities of SC utilization in Pakistan.

**Objectives:** To assess the level of knowledge, perception, and attitude of medical students regarding stem cell research (SCR) and its application, to obtain a better insight into the future of stem cell therapy in Pakistan as it is a rapidly emerging field in medicine.

**Materials and Methods:** This cross-sectional study was carried out using a self-administered questionnaire filled by 206 medical students from different medical colleges in Pakistan. A convenience sampling method was used. Knowledge and attitude scores were calculated based on answers to 28 well-structured questions. Data was analyzed using SPSS.

**Results:** The mean values of the answers showed that 60.2% (n=124) of the students had a good knowledge and 39.8% (n=82) had poor knowledge of stem cells. Whereas, 56.8% (n=117) expressed a positive attitude and 43.2% (n=89) expressed a negative attitude towards SCR. Independent t-test applied on knowledge score and attitude showed that the mean knowledge score of people with a positive attitude is higher i.e. 21.25 as compared to the mean knowledge score of people with negative attitude i.e. 19.21. And the difference of the means is significant at p=0.007. Thus, the attitude of students was observed to be significantly dependent on their knowledge about SCR.

**Conclusion:** The results show that medical students have baseline knowledge about SC therapy and a positive attitude towards it. Seminars, workshops should be conducted and this topic should be added to their syllabus so that they obtain proper information about SCR and encourage further research.

## INTRODUCTION

Stem cells (SC) are clonal cells that can differentiate into other types of cells and possess the property of self-renewal through mitotic divisions [1]. They are uncommitted progenitor cells, present in all multi-cellular organisms, which give rise to characteristic cells of organs and tissues. These cells are unique in their ability to keep on dividing and regenerate their population, in contrast to the differentiated cells, which do not divide and deplete if they are damaged. On the basis of potency, they are classified as uni-potent cells; which can differentiate into single mature cell type, pluripotent cells; which can give rise to most types of cells necessary for fetal development and totipotent cells; which can give rise to all cells types found in the fetus [2]. The sources to obtain these cells can be Placenta (Cord SCs), Fetal tissue or blastocyst (Embryonal SCs) and Blood, tissue or bone marrow (Adult SCs) [3].

Hematopoietic stem cell transplantation is an established treatment method for bone marrow failure diseases. The first allogeneic transplantation was performed by E. Donnall Thomas in 1957 and in Pakistan, the first transplant was done in 1995 at Dr. Ziauddin Hospital by Dr. Tahir Shamsi [4,5]. Recently, advances have been made towards the application of SC therapy for the treatment of diseases like Alzheimer’s, diabetes, immune-genetic conditions, cancers, Parkinson’s etc [6,7,8].

Research is being conducted increasingly in the field of cell biology worldwide in the light of its potential therapeutic benefit. It is becoming a popular option for treatment of those diseases that did not have adequate management available in the past. The use of stem cells has given birth to a new era of therapeutics which is known as regenerative medicine. Their renewal property offers exciting possibilities in reversing tissue damage caused by metabolic and degenerative changes. These scientific advancements require the healthcare workers to be equipped with knowledge regarding better innovative treatment options.

Guidelines for SC research in Pakistan have been developed by the National Bioethics Committee, Pakistan and adopted by the Human Organ Transplant Authority. However, it is still relatively new in Pakistan with less than twenty stem cell research institutes and limited awareness regarding the application of stem cell therapy among the healthcare workers, medical students as well as the general public. There is also a deficit in studies deducing the knowledge and attitude regarding stem cell research among the medical community in Pakistan. This demands avid exploration into the domain of stem cell research where the medical students lack knowledge. It indicates the significance of studying the cultural and religious factors that govern the level of interest and attitude of medical students regarding the subject. It is therefore timely that research be conducted to assess the level of knowledge, perception and attitude of medical students regarding stem cell research and its application to obtain a better insight into the future of SC therapy in Pakistan. Identification of the indistinct areas can better enable the medical community to take concrete measures to positively impact the students’ attitudes and supplement their knowledge about stem cells research. Better awareness can positively impact the progress in the field of regenerative medicine.

## Materials and Methods

This cross-sectional study was conducted from May 2019 to June 2019 after procuring an Approval Letter from the Ethical Review Board of the authors’ university. A convenience sampling method was used. A well-structured online questionnaire was made to be filled by 206 medical students from different medical colleges of Pakistan voluntarily. Full confidentiality was assured to the participants.

The questionnaire consisted of three sections; the Section A collected the socio-demographic data including age, sex, ethnicity, religion, institute and year of study. The religiosity and consideration of ethical aspect of SC research was also questioned.

The Section B included 18 questions regarding the students’ general and specific knowledge of stem cells. Score ‘2’ was set to the correct answer, score ‘1’ was set if the answer was ‘don’t know’ and score ‘0’ was set for the wrong answer. Thus the highest possible score for each student was 36. Mean of the total knowledge score(x=20) was taken as a cut-off value for good knowledge and poor knowledge. A score greater than 20 was considered as good knowledge while a score of 20 and below was considered as poor knowledge.

Section C comprised of 10 questions structured after literature review to assess the attitude of students towards stem cell research and therapy. Mean of x=32 was considered as a cut-off value for a positive attitude. A score of greater than 32 was considered as positive attitude and a score of 32 and below was considered as negative attitude.

Data was analyzed using SPSS version 23.0. To assess the internal consistency of the questionnaire the Cronbach Alpha coefficient was calculated to be 0.75. Descriptive statistics (frequency and percentages) identified demographic characteristics and students’ responses to the questionnaire. Paired t tests were used to analyze the relationships and statistical significance was considered to be as p value < 0.05. Pearson Test was used to find the correlation between knowledge and attitude of students towards stem cell research. Analysis included chi-square test to find significant association between knowledge and attitude of medical students towards stem cell research.

## Results

A total of 206 students from all five years of MBBS participated in this survey. The mean age of the respondents was 21 ± 1.43 years. Majority of them were females 76.7% (n=158) and 23.3% were males (n=48). All of them were Muslims. There were more responses from Year 3 and 4 as illustrated in Table 1.

**Table 1.** Demographic details of students.

|  | Students (n=206) |  | Frequency | Percentage |
| --- | --- | --- | --- | --- |
| **GENDER** | Gender | MALE | 48 | 23.3% |
|  |  | FEMALE | 158 | 76.7% |
| **YEAR OF STUDY** | Year Of Study | 1<sup>st</sup> Year | 25 | 12.1% |
|  |  | 2<sup>nd</sup> Year | 19 | 9.2% |
|  |  | 3<sup>rd</sup> Year | 44 | 21.4% |
|  |  | 4<sup>th</sup> Year | 87 | 42.2% |
|  |  | 5<sup>th</sup> Year | 31 | 15% |

**Table 2:** Assessing stem cell knowledge.

| # | Question | Correct | Incorrect | Don't know |
| --- | --- | --- | --- | --- |
| 1 | Do you have a generic knowledge of stem cells? | 90.8%(187) | 8.3%(17) | 1%(2) |
| 2 | Do you have a specific knowledge of stem cells? | 35.9%(74) | 58.7%(121) | 5.3%(11) |
| 3 | Did you ever attend vocational and training courses or meetings regarding stem cells? | 72.8%(150) | 25.2%(52) | 1.9%(4) |
| 4 | Would you be interested in developing your knowledge about stem cells? | 89.3%(184) | 8.7%(18) | 1.9%(4) |
| 1 | Do you know about the therapeutic applications with stem cells? | 66.5%(137) | 25.7% (53) | 7.8% (16) |
| 2 | Stem cells can be used to test new drugs. | 78.2%(161) | 7.8% (16) | 14.1% (29) |
| 3 | Stem cells can be used to treat Parkinson's, Alzheimer's, cancer, diabetes or heart diseases | 69.4%(143) | 13.1% (27) | 17.5% (36) |
| 4 | Stem cells can **divide and re-new** for long periods. | 92.2%(190) | 2.9% (6) | 4.9% (10) |
| 5 | Embryonic stem cells are un-specialized and capable of forming any cell type in the body including placenta. | 86.4%(178) | 7.8% (16) | 5.8% (12) |
| 6 | Stem cells can give rise to cancers. | 58.35%(120) | 22.3%(46) | 19.4%(40) |
| 7 | It is possible to isolate stem cells from human embryos (without their sacrifice) | 66.5%(137) | 13.1% (27) | 20.4% (42) |
| 8 | It is possible to isolate stem cells from the umbilical cord. | 56.3% (116) | 19.4% (40) | 24.3%(50) |
| 9 | Sperm and eggs are a source for adult stem cells. | 51% (105) | 30.6% (63) | 18.4% (38) |
| 10 | Adult stem cells are also known as somatic stem cells. | 65.5%(135) | 20.9%(43) | 13.6%(28) |
| 11 | Multipotent stem cells can be induced from normal skin cells by genetic reprogramming. | 58.7%(121) | 19.9%(41) | 22.3%(46) |
| 12 | Bone marrow stem cells are taken from the spine. | 50%(100) | 31.3%(64) | 18.9%(39) |
| 13 | Umbilical cord blood stem cell transplantation is less efficient compared to bone marrow stem cell transplantation. | 46.1%(95) | 21.4%(44) | 32.5%(67) |
| 14 | Stem cells are maintained by obligatory asymmetric replication i.e. one cell is similar to mother cell while other is a completely differentiated cell. | 40.3%(83) | 31.3%(64) | 28.6%(59) |

**Table 3.** Assessing attitude regarding stem cell research.

| # | Questions | Strongly disagree | Disagree | Not sure | Agree | Strongly agree |
| --- | --- | --- | --- | --- | --- | --- |
| 1 | I approve of stem cell research. | 8.7% (18) | 4.4%(9) | 17.5% | 51%(105) | 18%(37) |
|  |  |  |  | (36) |  |  |
| **2** | I am worried that stem cell transplantation might potentially open doors to human being killed for the benefit of others. | 4.4% (9) | 24.8%(51) | 33% (68) | 27.2% (56) | 10.2% (21) |
| **3** | The government should prohibit all researches regarding embryonic stem cells from embryo or aborted fetus. | 2.9%(6) | 8.3%(17) | 33.5% (69) | 35.4% (73) | 19.4% (40) |
| **4** | Life begins at conception; thus, embryonic stem cell research which involves the destruction of embryo is immoral, illegal and unnecessary. | 18%(37) | 23.3%(48) | 28.6% (59) | 16% (33) | 13.6%(28) |
| **5** | A blastocyst should be given the same respect and right to live as a living human adult. | 20.9%(43) | 33%(68) | 21.8% (45) | 11.2% (23) | 12.6%(26) |
| **6** | There IS A MORAL DIFFERENCE between creating embryos specifically for research and using embryos remaining after IVF (in vitro fertilization) for research? | 8.7% (18) | 11.7%(24) | 26.7% (55) | 37.9% (78) | 14.6% (30) |
| **7** | Embryos can specifically be made to be used for stem cell research and be destroyed during the study. | 17.5% (36) | 21.4% (44) | 32% (66) | 22.3% (46) | 6.3% (13) |
| **8** | Stem cell transplantation should be widely practiced. | 8.7% (18) | 13.1% (27) | 38.8% (80) | 31.6% (65) | 7.3% (15) |
| **9** | There should be awareness programs regarding stem cells and its use in research. | 8.7% (18) | 4.9% (10) | 21.8% (45) | 42.7% (88) | 21.8% (44) |
| **10** | The future of mankind is bright if stem cell research could be successfully conducted. | 8.7% (18) | 4.9% (10) | 21.8% (45) | 42.7% (88) | 21.8% (44) |

**Table 4.** Independent samples t-test results of knowledge and attitude.

|  | ATTITUDE | N | Mean | t | df | p |
| --- | --- | --- | --- | --- | --- | --- |
| **Knowledge Scores** | Positive | 117 | 21.25 | -2.720 | 204 | **0.007** |
|  | Negative | 89 | 19.21 |  |  |  |

Religiosity was self-reported by the respondents and most of them (76.2%) characterized themselves to as moderately religious.

**Fig 1.**
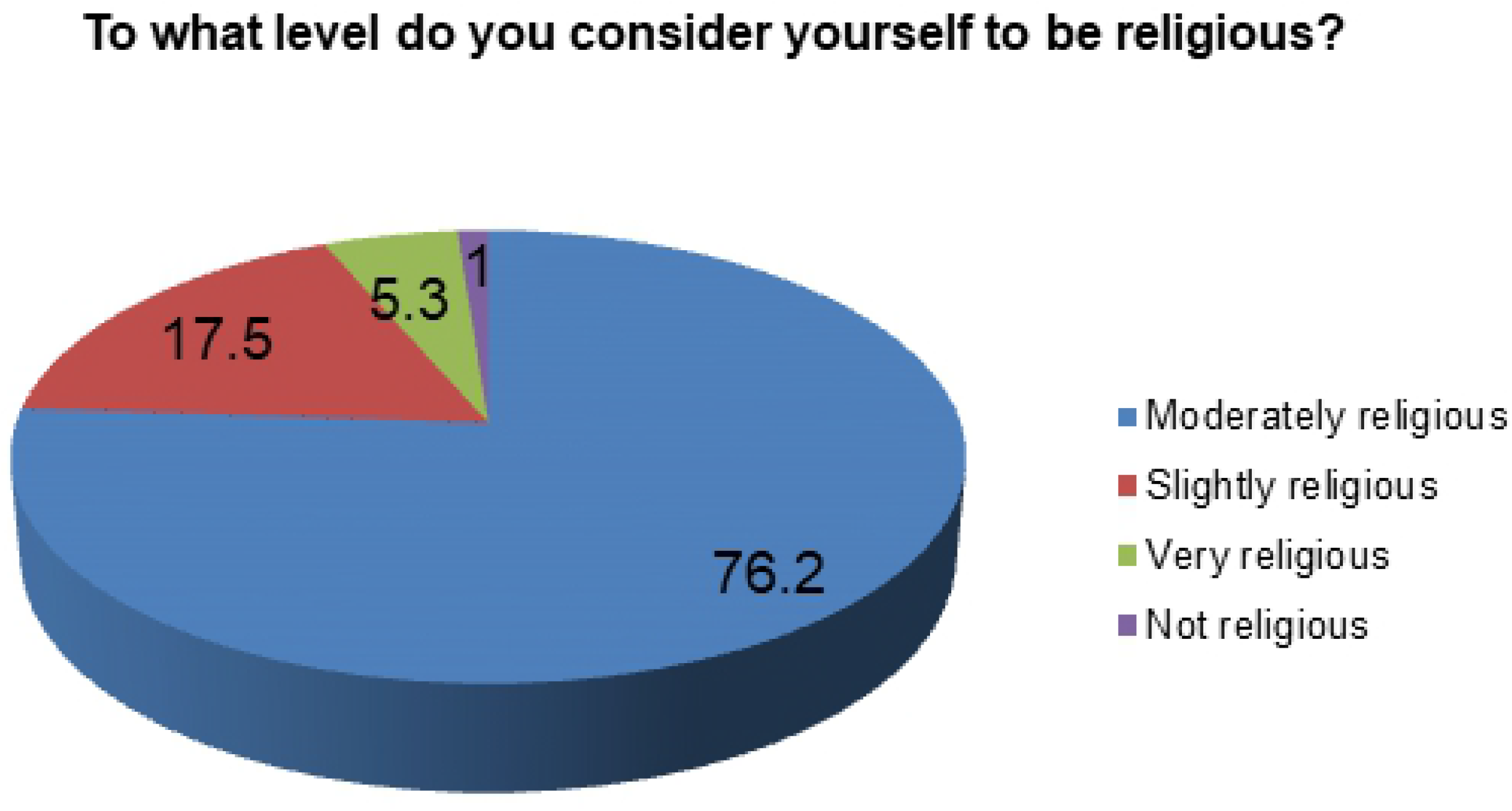
Self-reported religiosity.

They were further inquired whether they considered stem cell research to be ethical and 66% of the respondents did agree that SCR is ethical.

**Fig 2.**
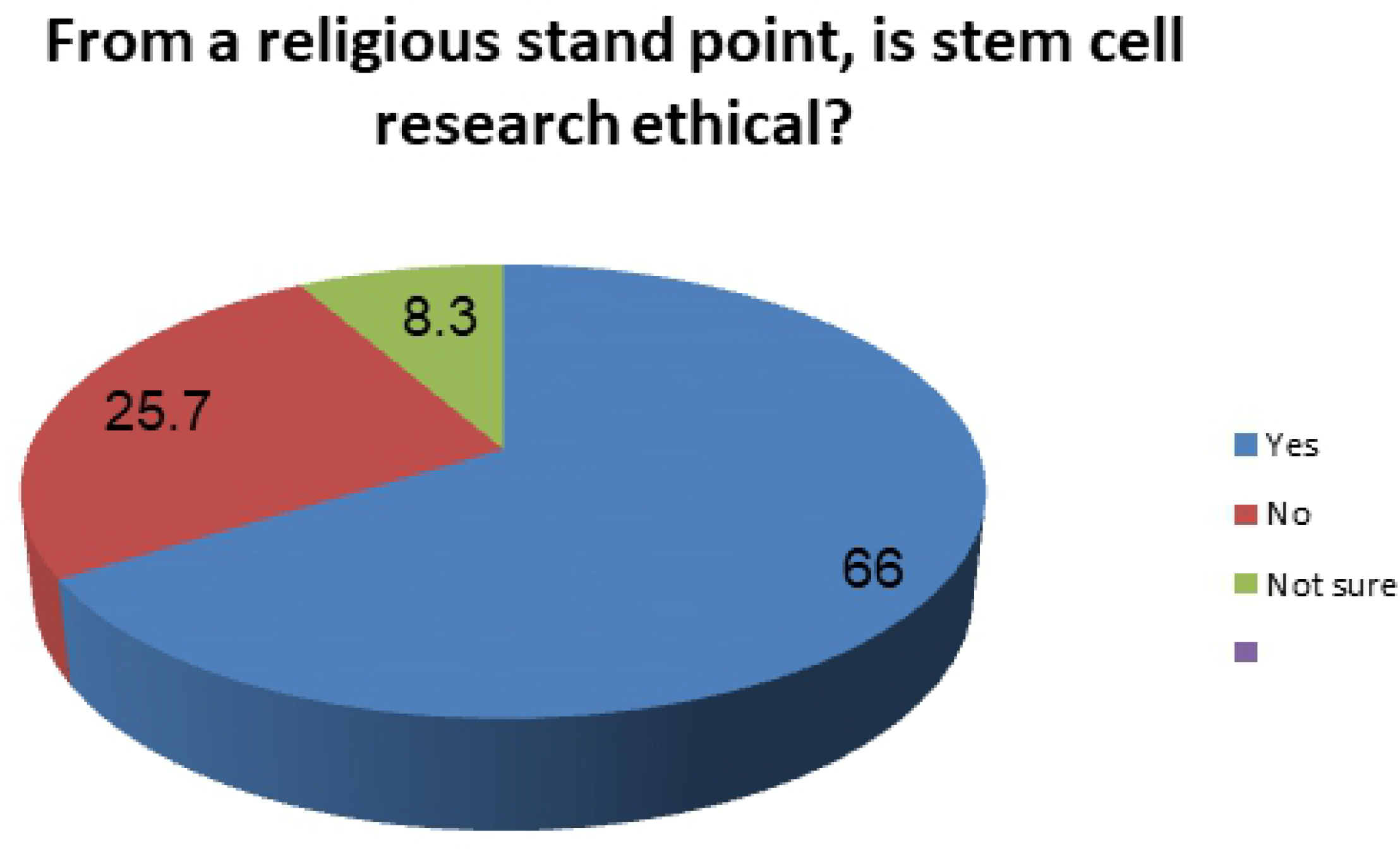
Ethical consideration.

Results showed that 90.8% (n=187) of respondents declared to have generic knowledge regarding stem cells but only 35.9% (n=74) had specific knowledge of stem cells. It was seen that 25.2% (n=52) had not learned about stem cells from vocational or training courses however, 89.3% (n=184) responded positively about interest in developing their knowledge about stem cells. For further evaluation of the students’ level of knowledge on stem cells, they were given a few questions and statements to answer.

Most of the students were aware of umbilical cord, embryonic and adult stem cells and the therapeutic applications of stem cells. However, it should be noted that only 2 participants answered correctly to all 14 questions of knowledge regarding stem cells.

The mean of knowledge scores was calculated as 20. All scores above 20 were considered to be good knowledge whereas a score of 20 and below was considered to be poor knowledge. Hence, out of n=206 students, 60.2% (n=124) were found to have good knowledge and 39.8% (n=82) had poor knowledge of stem cells.

**Fig 3.**
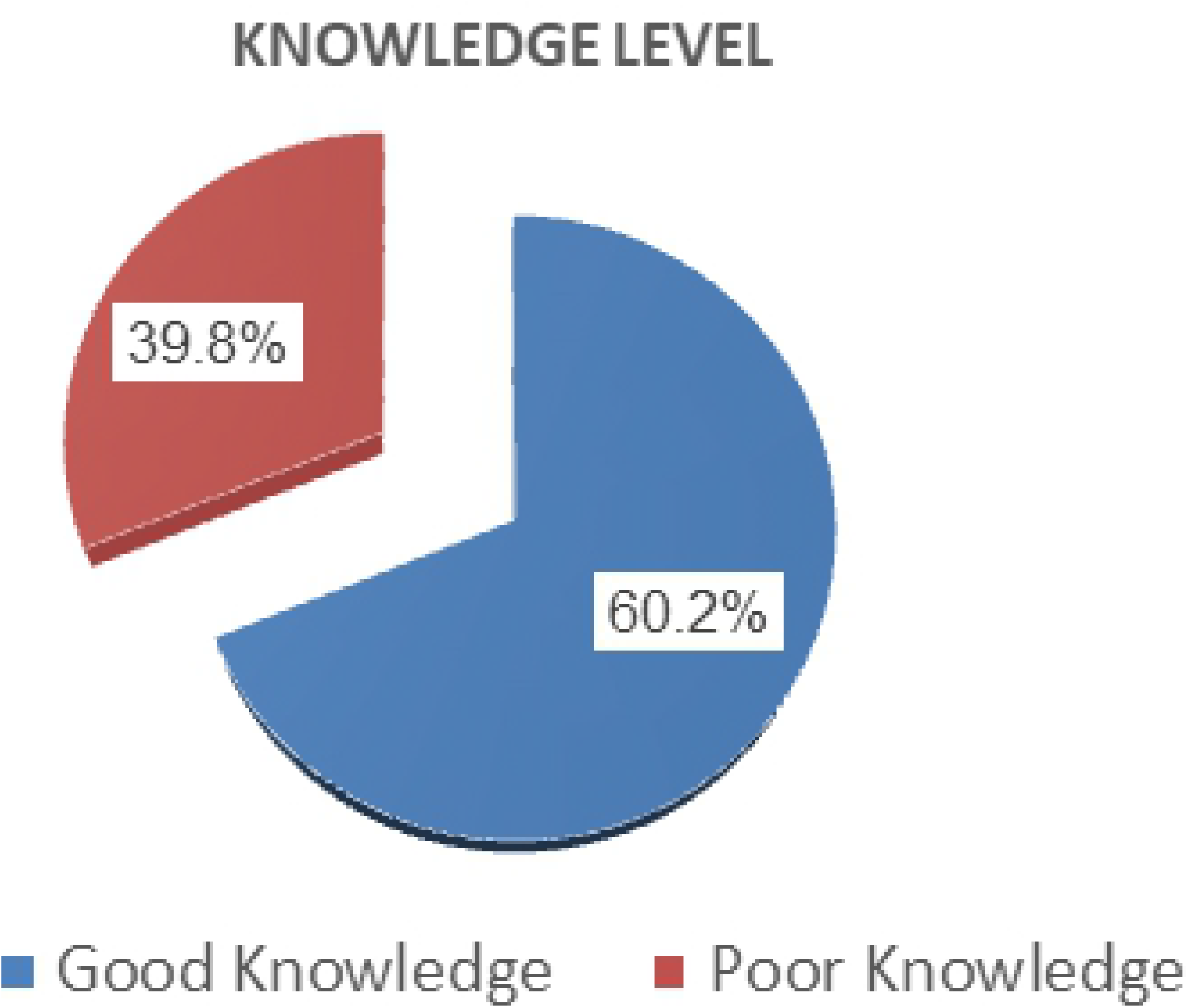
Result of knowledge scores.

To study the attitude and perception of students towards stem cell research, they were given a list of statements to rank their views on a 5 point Likert scale.

There were 52.5% (n=108) students who were of the opinion that there is a moral difference between creating embryos and using them specifically for research and using embryos remaining after In Vitro Fertilization (IVF) for research. Whereas only 29.6% (n=61) respondents believed that life begins at conception; thus, embryonic stem cell research which involves the destruction of embryo is immoral. There were also concerns by some students (n=77) about the potential of misuse of stem cell research for commercial purposes benefitting others.

The mean of the answers was taken and was calculated to be 32. Scores above 32 were considered to be a positive attitude and a score of 32 or below was considered to be a negative attitude. Results showed that 56.8% (n=117) expressed positive attitude and 43.2% (n=89) expressed negative attitude towards stem cell research.

**Fig 4.**
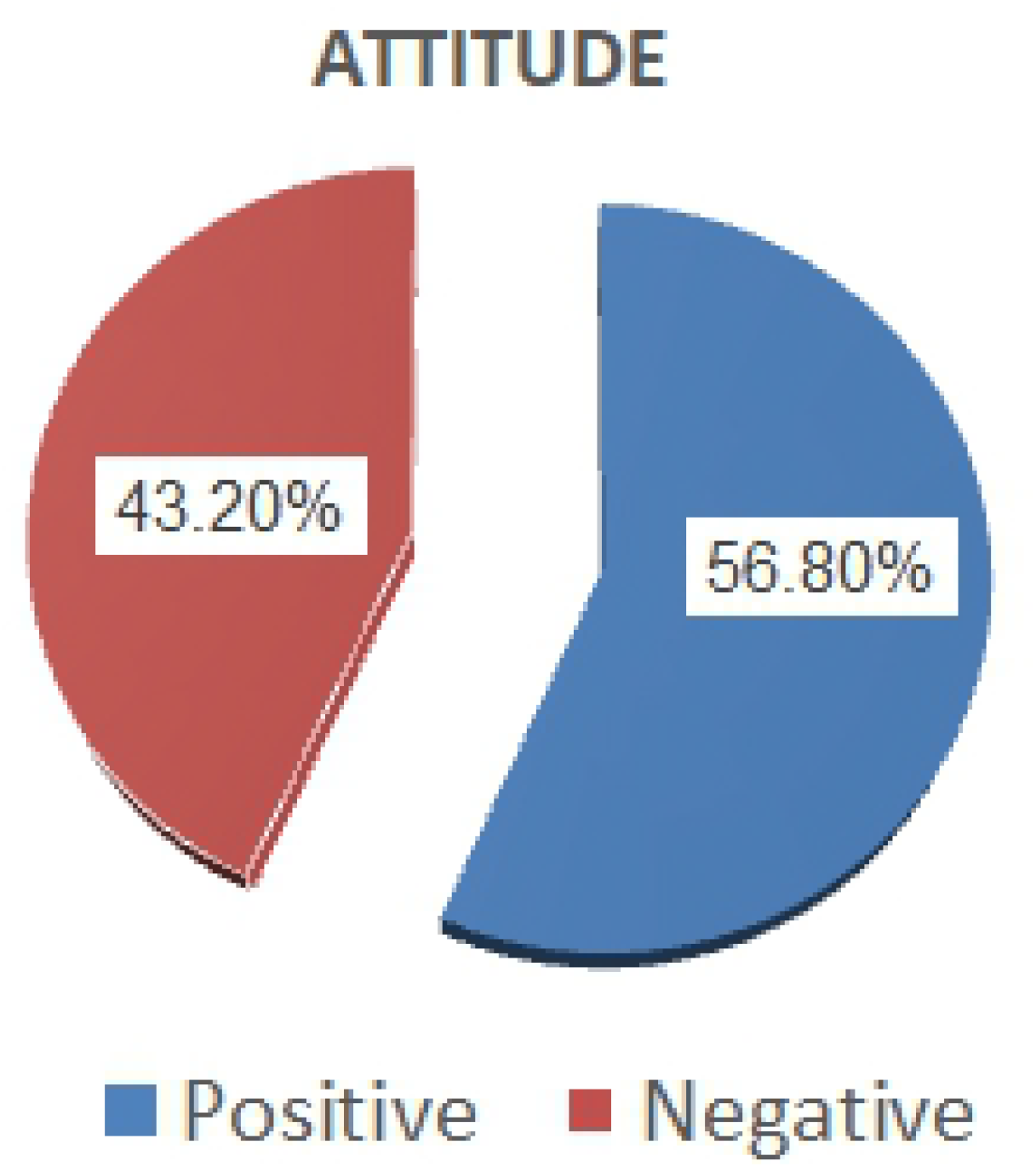
Result of attitude scores.

Independent t-test applied on knowledge score and attitude showed that the mean knowledge score of people with positive attitude is higher i.e. 21.25 as compared to the mean knowledge score of people with negative attitude i.e. 19.21. And the difference of the means is significant at p=0.007. Thus, increasing the awareness regarding stem cell research can increase its acceptability.

## Discussion

Stem cells have tremendous potential as it is clearly evident by the use of blood stem cells to treat diseases like leukemia; and can also be appreciated in the use of stem cells for tissue grafts to treat diseases or injuries to the bone, skin and eye [9].

In Pakistan, not much data is available on stem cell research with less than 100 articles published in PubMed. A stem cell society; Pakistan Stem Cells Society (PSCS) was established as late as 2012 [10]. Higher Education Commission (HEC) and Pakistan Science foundation (PSF) have approved many research projects on stem cells recently but they show slow development, mainly because of lack of funding, specific equipment and trained manpower.

According to our results, the knowledge level of the students is encouraging and suggests that they would be a trusted source of information, which would enable patients to make an informed decision regarding use of stem cells as a recent innovation in treatment. Our study showed a good knowledge level of 60.2% as compared to a similar study in KSA showing 31.2% good knowledge and 56% moderate knowledge among the respondents [11]. There was no association of stem cell awareness with gender, nationality, race, or year of study of the respondents in medical college corresponding to another study in Pakistan [12].

A strong majority of students (89.3%) also expressed willingness to develop their knowledge of stem cells in addition to the knowledge imparted through their curriculum which indicates that they are not neglectful of the topic and are encouraged to know more about it. Khali also reported that 92% of the respondents desired to attain coherent education about stem cells in a study conducted in Egypt [13].

It is also important to consider that stem cell research poses serious ethical and legal concerns and requires great responsibility especially when it comes to embryonic stem cells which can be created in laboratories solely for research. Majority of the respondents approved of stem cell research and believed it should be practiced widely. When considering from the religious point of view, it was seen that 66% students agreed that stem cell research is ethical. This majority is encouraging considering that all the students were Muslims and most of them considered themselves to be moderately religious. This is contrary to the findings of a study conducted in Australia where the Christian community considered embryonal stem cell research to be unacceptable [14].

Moreover, the distinction between creating stem cell specifically for research and using the surplus stem cells remaining after IVF was clearly understood by 52.5% of the students. This result correlated to a survey carried out in the US and Canada in 2008 where it was seen that 92% respondents supported stem cells derived from IVF as opposed to cloned embryos [15].

The students expressed concern over the misuse of stem cells research for promoting killing of human embryos for commercial purposes. This finding is in accordance to a study conducted in Greece where 73.6% were concerned that the umbilical cord blood could be used for purposes different than welfare in regenerative medicine [16].

## Conclusion

In the light of the findings of this study, it is concluded that the medical students of Pakistan showed statistically significant and affirmative knowledge as well as attitude towards stem cell therapy which indicates that in the near future, the field of applied biomedical sciences will show progress in leaps and bounds comparable to international standards. It is also encouraging to note that religiosity does not pose a significant threat to the future of SCR in Pakistan.

## Recommendations

Stem cell therapy heralds a new dawn in the treatment of many prevalent diseases which can be very favorable for mankind.

- The topic of stem cells and its therapeutic prospects must be made an extensive part of the medical curriculum to fill the theoretical knowledge deficits and inspire research in this field among students.
- The government should provide adequate funds for projects to promote both basic and applied stem cell research.
- Further studies should involve a larger sample population involving all medical colleges in Pakistan so as to obtain a more generalized conclusion.
- There should be seminars, symposiums and workshops in research centers and hospitals so that information and expertise could be shared and interest is developed in our young researchers and clinicians.

## Limitations

The limitations should also be taken into account for the results that have been presented. The small sample size presents a limitation and therefore the results may not be generalizable.

## Acknowledgments

The authors wish to thank the students of Rawalpindi Medical University for their cooperation.

## References

1) Weissman IL, Anderson DJ, Gage F. Stem and Progenitor Cells: Origins, Phenotypes, Lineage Commitments, and Transdifferentiations. Annu Rev Cell Dev Biol. 2001;17:387–403.

2) Sivakumar M, Dineshshankar J, Sunil P, Nirmal R, Sathiyajeeva J, Saravanan B. Stem cells: An insight into the therapeutic aspects from medical and dental perspectives. Journal of Pharmacy and Bioallied Sciences. 2015;7(6):361.

3) Arey L Developmental Anatomy: A Textbook and Laboratory Manual of Embryology. Ind Med Gaz. 1947 Jul;82(7):434–5.

4) Henig I, Zuckerman T. Hematopoietic stem cell transplantation—50 years of evolution and future perspectives. Rambam Maimonides medical journal. 2014 Oct;5(4).

5) Shamsi TS, Hashmi K, Adil S, Ahmad P, Irfan M, Raza S, Masood N, Shaikh U, Satti T, Farzana T, Ansari S. The stem cell transplant program in Pakistan—the first decade. Bone marrow transplantation. 2008 Aug;42(1):S114–7.

6) Dylla SJ, Beviglia L, Park IK, Chartier C, Raval J, Ngan L, Pickell K, Aguilar J, Lazetic S, Smith-Berdan S, Clarke MF. Colorectal cancer stem cells are enriched in xenogeneic tumors following chemotherapy. PloS one. 2008;3(6).

7) Damme A, Driessche T, Collen D, Chuah MK. Bone marrow stromal cells as targets for gene therapy. Current gene therapy. 2002 May 1;2(2):195–209.

8) Das S, Tobe B, Jain PA, Niles W, Winquist A, Mastrangelo L, Snyder EY. The Application and Future of Neural Stem Cells in Regenerative Medicine. InTranslational Regenerative Medicine 2015 Jan 1 (pp. 403–413).

9) Wert GD, Mummery C. Human embryonic stem cells: research, ethics and policy. Human reproduction. 2003 Apr 1;18(4):672–82.

10) Zahra SA, Muzavir SR, Ashraf S, Ahmad A. Stem cell research in pakistan; past, present and future. International journal of stem cells. 2015 May;8(1):1.

11) Tork H, Alraffaa S, Almutairi KJ, Alshammari NE, Alharbi AA, Alonzi AM. Stem cells: knowledge and attitude among health care providers in Qassim region, KSA. International Journal of Advanced Nursing Studies. 2018;7(1):1–7.

12) Khalil AM, Sharshor SM. Pediatric Nurses Knowledge, Awareness and Attitude towards Application of Stem Cells Therapy in Children. IOSR Journal of Nursing and Health Science. 2016;5(4):88–96.#

13) Mohammed HS, El Sayed HA. Knowledge and attitude of maternity nurses regarding cord blood collection and stem cells: An educational intervention. Journal of Nursing Education and Practice. 2015 Apr 1;5(4):58.

14) Tuch BE. Stem cells: a clinical update. Australian family physician. 2006 Sep;35(9):719.

15) Downey R, Geransar R. Stem cell research, publics’ and stakeholder views. Health Law Review. 2008;16(2):69.

16) Hatzistilli H, Zissimopoulou O, Galanis P, Siskou O, Prezerakos P, Zissimopoulos A, Kaitelidou D. Health Professionals’ knowledge and attitude towards the Umbilical Cord Blood donation in Greece. Hippokratia. 2014 Apr;18(2):110.

